# Recombinant protein condensation inside E. coli enables the development of building blocks for bioinspired materials engineering – biomimetic spider silk protein as a case study

**DOI:** 10.1101/2022.09.14.507915

**Authors:** Bartosz Gabryelczyk, Fred-Eric Sammalisto, Julie-Anne Gandier, Jianhui Feng, Grégory Beaune, Jaakko V. I. Timonen, Markus Linder

## Abstract

Recombinant expression of proteins destined to form biological materials often results in poor production yields or loss of their function due to premature aggregation. Recently, liquid-liquid phase separation has been proposed as a mechanism to control protein solubility during expression and accumulation in the cytoplasm. Here, we investigate this process *in vivo* during the recombinant overexpression of the spider silk-mimetic mini-spidroin NT2RepCT in *Escherichia coli*. The protein forms intracellular liquid-like condensates that shift to a solid-like state triggered by a decrease in their microenvironmental pH. These features are also maintained in the purified sample *in vitro* both in the presence of a molecular crowding agent mimicking the bacterial intracellular environment, and during a biomimetic extrusion process leading to fiber formation. Overall, we demonstrate that characterization of protein condensates inside *E. coli* could be used as a basis for selecting proteins for both materials applications and their fundamental structure-function studies.

## 1. Introduction

Liquid-liquid phase separation (LLPS) is a ubiquitous process found throughout biology as a means of controlling the local concentrations of proteins or RNA [1, 2]. LLPS leads to the formation of a condensed biomolecule phase —also called condensate— observed usually as micron-size spherical droplets. Intracellularly LLPS is involved in the formation of membraneless organelles associated with regulation of various cellular processes [3]. Extracellularly, protein LLPS has been shown as a key first step in the assembly pathways of various biological materials [4], for example elastomeric structures made of elastin or resilin [5, 6], underwater adhesives secreted by marine organisms [7, 8], protein-based composites [9, 10], and spider silk-like fibers [11, 12].

LLPS has gained significant attention in recent years and the features of proteins that drive biological material assembly are increasingly understood. For example, it has been suggested that LLPS is used by natural systems to control the physical state of protein assembles, i.e. their material state. LLPS provides a way to regulate proteins’ transition from a soluble (liquid-like state) into the solid final form of a material. During this process the condensed phase can also serve to prevent premature aggregation [13, 14].Therefore, protein LLPS has emerged as a new concept in bioinspired material engineering. In this approach proteins derived from sequences identified in biological materials are recombinantly produced, purified and assembled through LLPS into bioinspired materials in a biomimetic process. However, protein engineering strategies that put into practice these new assembly pathways in material applications lag in comparison. This can be attributed to the challenges associated with the recombinant expression of proteins derived from biological structures. In their native forms, these proteins often contain intrinsically disordered regions, long and repetitive sequences and/or amyloidogenic motifs. When such native sequences are recombinantly overexpressed, in most cases, they exhibit very low production yield or are prone to premature aggregation and accumulation into insoluble inclusion bodies [15, 16, 17, 18]. Protein sequestration into inclusion bodies during recombinant expression in bacteria is a widely used strategy to achieve very high production yield [19, 20]. However, aggregated proteins that accumulate in the inclusion bodies often require re-solubilization and refolding. Sometimes harsh conditions and chemicals need to be used which not only add expenses and steps to the process, but can also result in the modification or fragmentation of the proteins [21, 22]. Solubilization can also affect subsequent processing and assembly of the proteins and therefore is not necessarily compatible with the biomimetic principles governing the fabrication of protein-based materials [23].

Recently, an engineered spider silk protein i.e. mini-spidroin (NT2RepCT) was developed [24]. The protein is composed of the globular N-terminal (NT) and C-terminal (CT) domains present in the native spidroins and are linked by two repetitive poly-alanine blocks (2Rep) derived from the intrinsically disordered central domain of spidroins. The terminal domains regulate solubility of silk proteins and mediate their assembly into a solid fiber, initiated by a pH-dependent dimerization of the NT-domains [25, 26, 27]. In the protein’s native form, the repetitive region linking these domains is composed of approximately 100 modular units and is associated with the mechanical properties of the silk fibers [28]. It was found that shortening of this aggregation-prone central repetitive region to only two units enables high recombinant expression yields in *E. coli* of up to 20 g/L [29]. Moreover, the presence of the NT domain in the protein sequence enables an extreme solubility in an aqueous buffer at pH 8 after purification and additionally allows assembly into fibers using biomimetic spinning setups [24, 29, 30].

In this study, we investigated whether NT2RepCT undergoes LLPS and if the formation of protein condensates can be linked to its high expression yield and ability to form silk fibers. We studied LLPS properties of the protein form the beginning of its overexpression and accumulation within *E. coli* to the extrusion of the purified protein during fiber formation in a biomimetic silk spinning process. Thus, we begin to establish the critical connections between intracellular phase behavior of the protein *in vivo* with its ability to form functional materials *in vitro*.

## 2. Methods

### 2.1. Cloning

DNA sequences encoding eGFP-NT2RepCT, NT2RepCT, and eGFP (with N-terminal 6xHis-tags appended to each) were inserted into a pET-28a expression vector using golden gate cloning [31]. The DNA sequence of this expression vector is the same as its commercially available pET-28a counterpart with the exception of the insertion of two recognition sites for the BsaI restriction enzyme in its multiple cloning site. This modification enabled its use with the golden gate method. The KSI-eGFP was obtained from Genscript (prepared by inserting the synthesized eGFP gene after KSI sequence that was already present in pET-31b(+) expression vector, using XhoI/AlwNI cloning sites).

### 2.2. Protein expression

All plasmids were transformed into strain BL21AI *E. coli* (obtained from Thermo Fisher Scientific). *E. coli* cells were cultivated in LB medium (volume = 3 mL) containing an appropriate antibiotic, overnight at +30 °C at 220 RPM. Next, the cultures were diluted 1/100 with the growth medium and incubated in the same conditions until OD_600_ reached 0.6. Then the temperature was lowered to +20 °C, the protein expression was induced with 0.2 % L-arabinose and 0.5 mM IPTG and carried out for 18 h. Expression of NT2RepCT and eGFP-NT2RepCT for *in vitro* studies was carried out in 500 mL culture volume using the same protocol as described above. Cells were harvested by centrifugation (5000 RPM, +4 °C, 10 min). Cell pellets were frozen at -20 °C.

### 2.3. Purification of NT2RepCT and eGFP-NT2RepCT

Frozen cells were thawed and resuspended in 20 mM Tris-HCl buffer pH 8, containing 15 mM imidazole. Cell lysis was performed using emulsiflex cell homogenizer (18000 psi, +4 °C) after which the lysate was centrifuged at 20,000 PRM at +4 °C for 20 min. The soluble fraction was loaded onto a 5 mL Ni-NTA column connected to an ÄKTA pure (GE Healthcare) chromatography system. The target protein was eluted from the column with 20 mM Tris-HCl pH 8 containing 300 mM imidazole. The eluted protein was dialyzed against 20 mM Tris-HCl pH 8, at +4 °C using a SnakeSkin dialysis membrane (Thermo Fisher Scientific) with a 3.5 kDa molecular-weight cutoff. Protein purity and integrity was verified with SDS-PAGE.

### 2.4. Fluorescence microscopy

Fluorescence and phase contrast images of bacterial cells were acquired using an Axio Observer Z1 microscope (Carl Zeiss, Germany) equipped with 100x/1.4 oil objective, 1.6x tube lens, and Andor iXon Ultra 888 camera. Bacterial cells were deposited onto standard microscope slides coated with poly-D-Lysine (A3890401 Thermo Fisher Scientific). eGFP signal was obtained using excitation light at 470 nm, while collecting the emitted light of 500 - 554 nm. Images of the cells at various time points (Figure 1) were acquired using different settings (exposure time and light source intensity).

**Figure 1.**
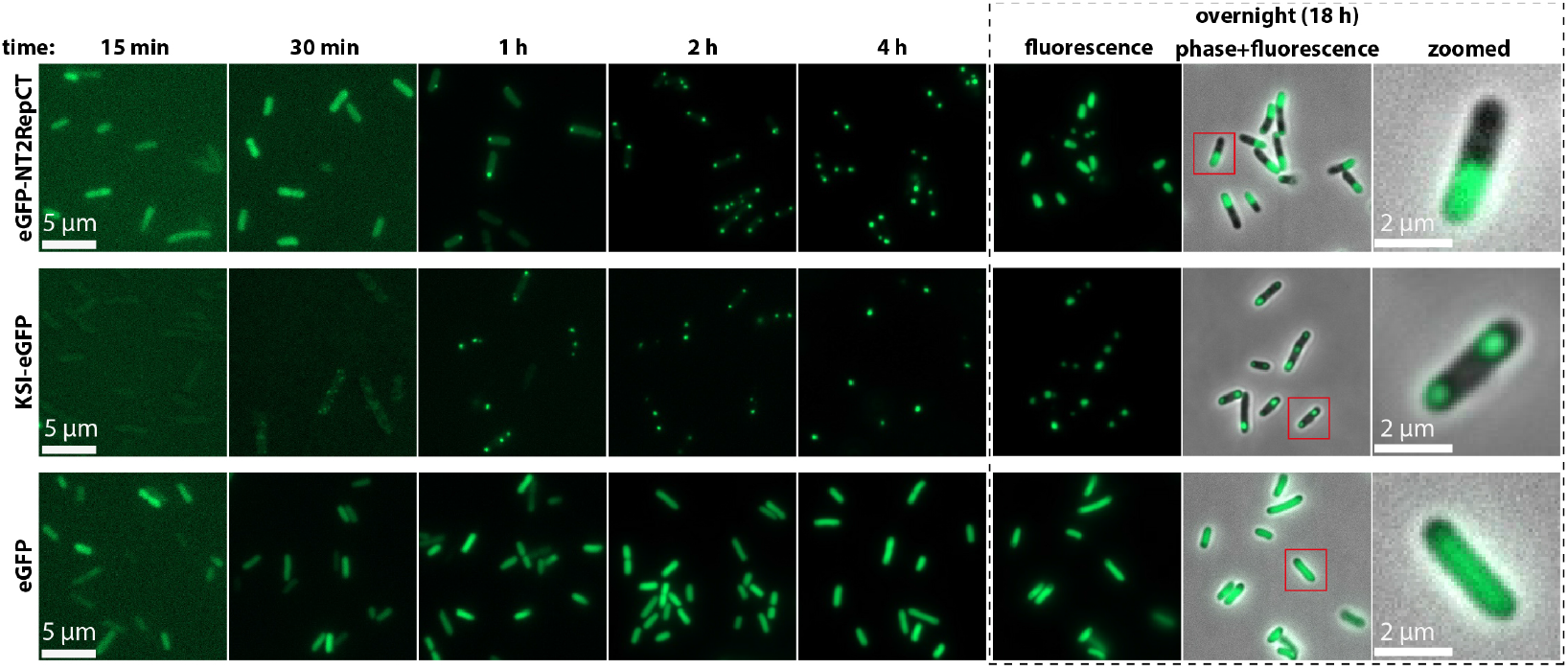
Fluorescence microscopy of *E. coli* cells expressing eGFP-NT2RepCT and the reference proteins eGFP and KSI-eGFP at different time points after induction of protein expression (t = 0 h). At the final time point (t = 18 h), fluorescence and phase contrast images are overlaid and shown at higher magnification (panel with dashed box). While differences between intracellular structures formed by each protein can be observed over time at low magnification, these differences are most prominent at high magnification at the final time point: eGFP without its fusion partner is uniformly spread, eGFP-NT2RepCT forms a distinct region occupying approximately half of the cell’s volume, and KSI-eGFP concentrates in two distinct spots at both poles of the cell.

### 2.5. Phase contrast microscopy

Images of protein samples during *in vitro* studies were acquired using an Axio Observer Z1 microscope (Carl Zeiss, Germany) equipped with 40x magnification objective and Axiocam 503 camera. Samples were prepared by mixing the protein (in 20 mM Tris-Cl pH 8.0) with an appropriate buffer on a glass slide in 1/5 (protein/buffer) volume ratio or 1/1 ratio (Figure 5c-e). For testing the influence of pH on the phase transition of the protein the following buffers were used: Tris-HCl (pH 7.5, 7), sodium phosphate (pH 6.5, 6.0), sodium acetate (pH 5.5, 5.0). The final concentration of each buffer was 100 mM. For the experiments mimicking the bacterial intracellular environment (Figure 5a and Figure S3a) the buffers contained an additional 10 % (w/v) of dextran 500 (Amersham Biosciences).

### 2.6. Fluorescence recovery after photobleaching (FRAP)

Bacterial cells suspended in LB growth medium were deposited onto a microscope coverslip (coated with poly-D-Lysine (A3890401 Thermo Fisher Scientific)) 18 h after induction of protein expression. Images were acquired using Nikon Ti-E inverted microscope equipped with 60x/1.4 oil objective lens, 1.5x tube lens, Crest Optics X-light V3 spinning disk confocal head (operated in the widefield mode) and Hamamatsu Orca Flash 4.0 LT+ camera. The system was controlled by *μ*Manager (2.0) open-source microscopy software. eGFP was excited using continuous illumination with 470 nm light from LDI Laser Diode Illuminator (89 North) at 1 % power level. Emission was collected between 485-535 nm with exposure time of 5 ms, allowing frame rates up to 200 fps. Local spot-like photobleaching was achieved using Gataca Systems iLas 2 unit coupled to a 100 mW OBIS LX 405 nm laser operated at 4 % power level. The 405 nm laser was focused to a circular spot with a diameter of approximately 1 micron and the photobleaching exposure time was manually controlled to be approximately 900 ms. Processing of the images and calculation of the fluorescence recovery was carried out using Fiji (2.3.0) software. Normalized fluorescence intensity was calculated as:

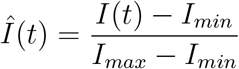

where *Î(t)* is the normalized fluorescence intensity, I(t) the measured fluorescence intensity (unprocessed 16-bit pixel values from the detector), and *I*_*min*_ and *I*_*max*_ are the minimum and maximum measured intensity values within the data set.

### 2.7. Flow cytometry

Bacterial cells were cultured for 18 h as described in the “Protein expression” section. Cells were then harvested by centrifugation 3000 RPM, 3 min, +20 °C and resuspended in 1x PBS buffer pH 7.4. Cell density was adjusted to 10^6^ cells/mL. Flow cytometry measurements were performed on a BD FACSAria II cytometer (BD Biosciences) using 488 nm laser for excitation and a 530/30 nm bandpass filter for emission detection. 10000 events per sample was recorded and the gated events were further analyzed. Data analysis was performed using the BD FACSDiva software package (version 6.0).

### 2.8. Partial cell lysis

Bacterial cells were harvested by centrifugation (3000 RPM, +20 °C, 3 min) 18 h after induction of protein expression. The cells were then gently suspended in 50 mM Tris-HCl pH 8.0 or 50 mM sodium phosphate pH 6.0. 1.25 – 1.5 *μ*l of cell suspension was pipetted between two microscope glass slides, pressed several times, and imaged with the fluorescence microscope.

### 2.9. Complete cell lysis

Bacterial cells were harvested by centrifugation (8000 RPM, +20 °C, 5 min) 18 h after induction of protein expression. Cell pellets were frozen at -20 °C, thawed, resuspended in B-PER Bacterial Protein Extraction Reagent (Thermo Fisher Scientific) containing 0.5 mg/mL of lysozyme (Sigma-Aldrich L7651), and then mixed with 100 mM Tris-HCl pH 8.0 or 100 mM sodium phosphate pH 6.0 in 1/1 volume ratio. Cell lysis was carried out for 10 min. 1.5 *μ*l of cell suspension was used for imaging with the fluorescence microscope. Remaining sample was centrifuged at 20000 RPM +20 °C, 5 min. Obtained soluble and insoluble fraction (pellet of cell debris) was analyzed by SDS-PAGE using 4–20% Mini-PROTEAN TGX Precast Protein Gel (Bio-Rad) and Precision Plus Protein Dual Color Standard (Bio-Rad).

### 2.10. Investigation of protein phase transition during extrusion

Purified NT2RepCT (suspended in 20 mM Tris-HCl pH 8.0 buffer) was concentrated to approximately 100 mg/mL using a Vivaspin centrifugal concentrator (Sartorius) with 10 kDa molecular-weight cutoff.

Micropipettes were prepared by pulling borosilicate capillaries (WPI, 1 mm/0.5 mm outer/inner diameter) using a puller (PN-31, Narishige). The micropipettes were then sized to approximately 20 *μ*m in diameter and bent using a microforge (MF-900, Narishige) to keep them horizontal during the experiments. The micropipette was connected by a tube to a reservoir filled with 20 mM Tris-HCl pH 8.0 buffer and linked to a piezoelectric pressure controller (Elveflow). The pressure was controlled using the instrument’s software and allowed the tubing and the pipette to be filled with the buffer solution.

The concentrated protein was aspirated in the micropipette using negative pressure, followed by 20 mM Tris-HCl pH 8.0 buffer. This produced a concentration gradient of the protein across the length of the micropipette by diffusion of the high concentration protein solution into the pure buffer. The pipette was immersed in a medium-sized (60 mm in diameter) petri dish (serving as a coagulation bath and observation chamber) filled with 500 mM sodium acetate buffer pH 5 containing 200 mM NaCl. The mixture was then extruded under constant positive pressure of 500 Pa. The Petri dish, located on a motorized stage, was displaced at a constant speed during the experiment to avoid blockage of the capillary outlet by the extruded protein at high concentration.

The experiments were performed at room temperature (T = 23 ± 1 °C)and visualized with an inverted microscope (Nikon Eclipse Ti) equipped with a 20x objective and associated with a 1.5x magnification device. Bright-field images were recorded with a sCMOS camera (Zyla-4.2-CL10, Andor), at a time interval of 50 ms, that was operated using *μ*Manager (2.0) open-source microscopy software. The images were processed with Fiji (2.3.0) software.

## 3. Results and Discussion

### 3.1. NT2RepCT undergoes LLPS during overexpression in E. coli forming protein condensates that exhibit liquid-like properties

First, we investigated the material state of the NT2RepCT in the cytoplasm of *E. coli* during recombinant overexpression, that is, whether the protein undergoes LLPS and forms intracellular protein condensates. Enhanced green fluorescent protein (eGFP) was fused to the N-terminus of the protein to enable its visualization by fluorescence microscopy. Ketosteroid isomerase (KSI) fused with eGFP at its C-terminus was used as a reference because it is known to form insoluble inclusion bodies during overexpression under the conditions studied [32]. In addition, eGFP without any fusion partner was also included in the study since it was previously found that it is recombinantly expressed in soluble form [33, 34]. Thus, it served as a second reference point for the study of the biomimetic silk protein’s state. To minimize basal protein expression (promoter leaking) and to ensure a precise time-course study, the *E. coli* BL21-AI strain was used as its T7 RNA polymerase is tightly regulated by an arabinose-inducible promoter [35, 36]. The alternative BL21(DE3) strain with T7 RNA polymerase under control of a lacUV5 promoter was also tested but was not practical due to its leakiness.

After growth to mid-exponential phase, recombinant protein expression in *E. coli* was induced and imaged over time (Figure 1). For eGFP-NT2RepCT, within the first two time points (t = 15 min and t = 30 min), an evenly distributed fluorescence was detected across the cytoplasm. After 1 h the fluorescence signal began to localize in the polar regions of the rod-shaped cells. The overall intensity of the fluorescence signal increased over time and began to colocalize mainly at one end of the cell after 4 h. Ultimately, at the final time point (18 h), the fluorescent intracellular structures occupied up to approximately half of the cell’s cytoplasm. In contrast, during expression of eGFP, the fluorescence signal gradually increased over time, and remained homogenously dispersed throughout the cell. Expression of the known inclusion body-forming KSI-eGFP also displayed distinct features. Interestingly, the overall fluorescence signal detected in the cells was lower compared to eGFP-NT2RepCT and eGFP. Furthermore, this fluorescence could only be visualized after 30 min. At this time, the distribution of the fluorescence was uneven, and small concentrated regions could be observed within the cell. Then, after 1 hour, fluorescence could be seen accumulating at both poles of the cells. These two polar structures became more prominent after 2 hours and in morphology resembled the intracellular structures observed for the eGFP-NT2RepCT at the same time point. However, as protein production continued, the morphological differences between the intracellular structures formed by these two proteins became more visible. At the final time point, in most of the cells the two initially observed structures for eGFP-NT2RepCT merged into one, but in the case of KSI-eGFP, remained concentrated at the two poles of the cell. The merging into one structure is the expected behavior for LLPS since this minimizes the overall surface energy. That is, overall, the formation of larger droplets at the expense of smaller ones is favored [37, 1].

Although the morphologies of the structures formed by eGFP-NT2RepCT and KSI-eGFP were clearly distinct at the final time point, they could not be associated with an insoluble or condensed state exclusively based on these features. For instance, both inclusion bodies [38, 39] and liquid-like protein condensates [40, 14, 34] have been observed to locate to either one or both poles of the *E. coli* cell with similar morphologies. As such, we probed their material state using fluorescence recovery after photobleaching (FRAP). By focusing a 405 nm laser beam to approximately 1 micrometer (full-width at half maximum, FWHM) we achieved partial bleaching of the intracellular structures and assessed their viscoelastic properties by monitoring the fluorescence intensity changes over time in the regions of interest (ROI1 and ROI2). Within the photobleached ROI1 of eGFP-NT2RepCT structures 36 % of the fluorescence was recovered within approximately 20 s. This is clearly visualized in the microscopy images of Figure 2a. Recovery, however, was not complete, given the anticipated minor photobleaching that reduces the number of fluorescently active proteins within the cell. In the ROI2, located outside of the bleached area, depletion of fluorescence over time was observed further supporting that diffusion of the fluorescent protein within the condensate can be linked to its liquid-like viscoelastic properties. In contrast, bleaching of the intracellular structure formed by the KSI-eGFP showed no recovery of the fluorescence in the bleached area (ROI1) nor depletion of the fluorescence outside the bleached area (ROI2) (Figure 2b). These results show that KSI-eGFP within the intracellular structures was not able to diffuse, which is consistent with a solid-like material state. Finally, the FRAP analysis of the cells expressing eGFP without any fusion partner showed very fast fluorescence recovery (time below 1 s) of the bleached area (Figure 2c – ROI1). At the same time depletion in the fluorescence in the region outside the bleached spot was observed (Figure 2c – ROI2), that all together, indicated fast diffusion of the eGFP and a highly fluidic behavior within the cell. These results are consistent with the previously made observations in the literature showing that under similar expression conditions eGFP is soluble in the cytoplasm while KSI-eGFP forms insoluble inclusion bodies [32, 33].

**Figure 2.**
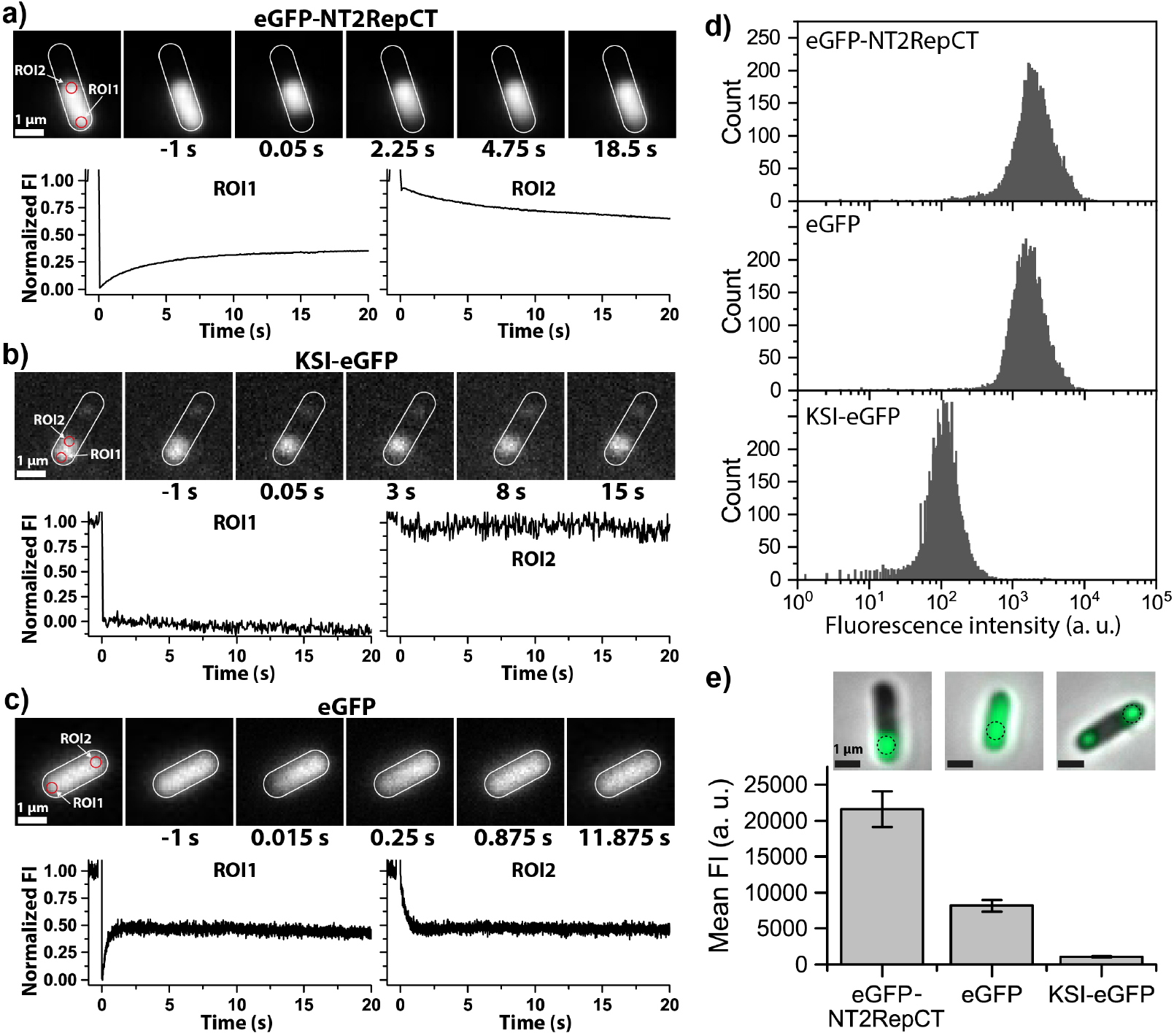
Investigation of the material state of intracellular structures formed by eGFP-NT2RepCT as well as reference proteins eGFP and KSI-eGFP 18 h after induction of protein expression. a-c) Fluorescence recovery after photo-bleaching (FRAP) – images of cells captured at different time points before (−1 s) and after photobleaching (0.05-18.5 s). The contour of the cell is approximately marked by the white line and the regions of interest (ROIs) used for recovery analysis marked with red circles. Plots below the images correspond to the normalized fluorescence intensity (FI) within the photobleached area (ROI1) and outside of the photobleached area (ROI2). Data normalization was carried out using equation (1) presented in the methods section. d) Flow cytometry – histograms show the differences between overall fluorescence intensity recorded for bacterial cells expressing the proteins under study e) Mean fluorescence intensity measured from the microscopic images at ROIs (dashed line) located within the intracellular structures observed. Plot shows average of 15 measurements acquired from different cells.

In addition to FRAP, we also used flow cytometry to analyze the fluorescence intensity of the bacterial cells at the final time point of the protein overexpression. We observed that the fluorescence of the cells expressing eGFP-NT2RepCT and eGFP was in the same range, while it was an order of magnitude lower for cells expressing KSI-eGFP. (Figure 2d) Therefore, we additionally analyzed the mean fluorescence signal intensity from the microscopic images (acquired using the same settings for all samples) measured from ROIs (approx. 0.5 *μ*m^2^) located within the intracellular structures Figure 2e – images). We observed that the fluorescence signal recorded from the eGFP-NT2RepCT had the highest intensity, that was approximately 2.5x higher than eGFP, and 20x higher compared to KSI-eGFP (Figure 2e – bar plot). These results indicated that eGFP-NT2RepCT was highly concentrated within the condensates. On the other hand, restricted molecular mobility, or protein misfolding as a result of aggregation might lead to reduced fluorescence of KSI-eGFP. These observations are in agreement with FRAP analysis and are further supported by the SDS-PAGE showing all the proteins had similar expression level at the final point of the protein expression (Figure S1). Therefore, the differences in fluorescence intensity of the intracellular structures formed by the proteins under study are in-line with their anticipated material state.

### 3.2. NT2RepCT intracellular condensates change their material state from liquid- to solid-like when exposed to pH 6

Next, we investigated whether a decrease in pH could trigger a transition of the liquid-like intracellular condensates of NT2RepCT into a solid-like state. We partially disrupted the bacterial cells by squeezing them between two microscope glass slides at either pH 8 or pH 6 (Figure 3). This treatment allowed us to observe in real time, the spreading of the fluorescent material into the extracellular space — indicating a liquid-like state of the protein — or lack thereof, indicating a solid-like state of the protein. As seen in Figure 3a, bacterial cells expressing eGFP-NT2RepCT displayed a pH dependent behavior as anticipated. At pH 8.0 partial cell damage led to spreading of the intracellular structures outside of the cells while at pH 6.0 the structures remained intact, and no spreading of the fluorescence was observed. This behavior suggests a shift of a material state of the protein structures from liquid-like to solid-like with a decrease in pH showing that the properties of the intracellular structures are consistent with the behavior of the purified NT2RepCT and native spidroins. Indeed, the pH-triggered transition of soluble NT2RepCT to solid state occurs also during silk fiber production in a biomimetic setup [24]. The same process occurs also during assembly into fibers of native spidroins in the spinning duct of a spider [26, 41]

**Figure 3.**
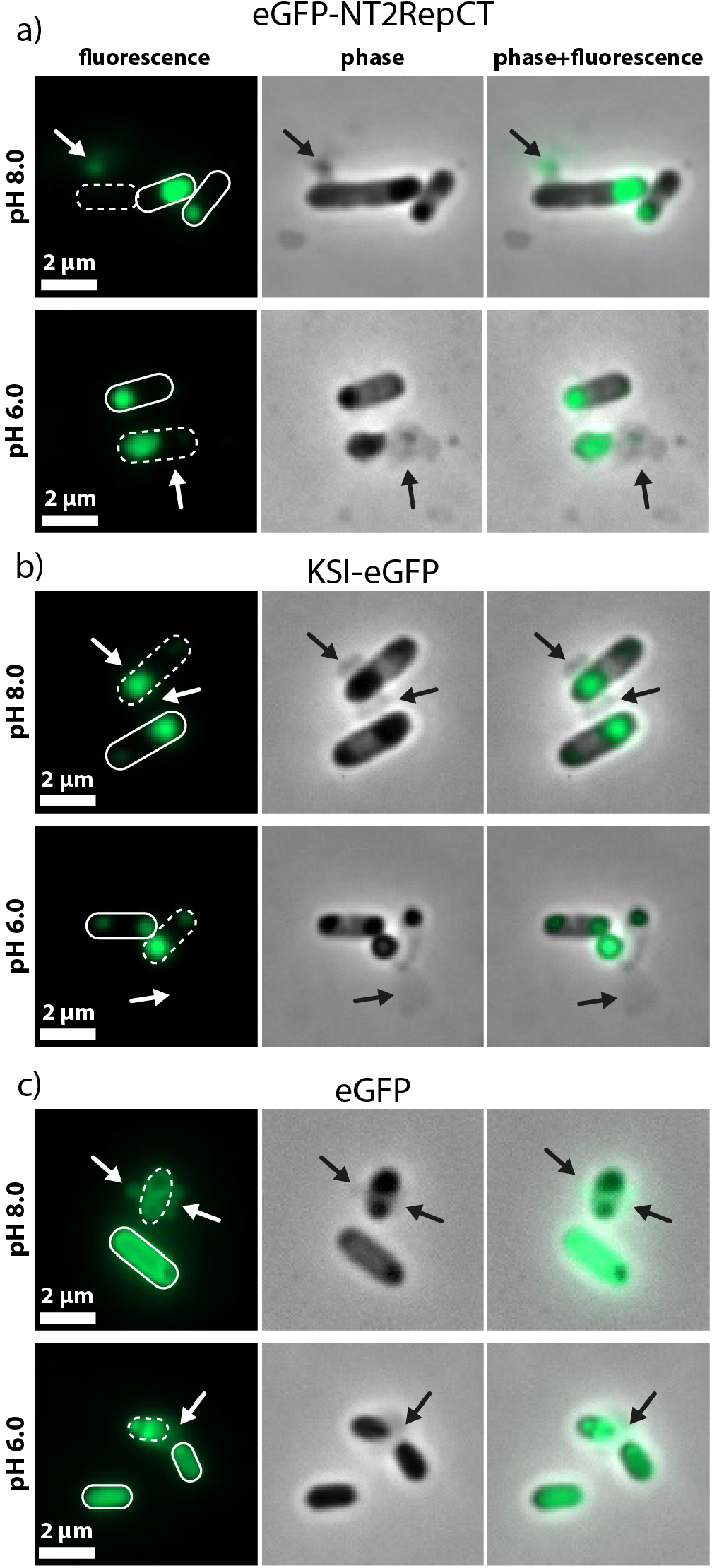
Fluorescence and phase contrast microscopy of intact and partially disrupted *E. coli* cells at pH 6.0 and pH 8.0. Damaged cells (by squeezing between two microscope glass slides) are indicated by a dashed line while intact cells are indicated by a solid line. Arrows show the spreading area of cellular content from disrupted cells. Spreading of the fluorescence indicates liquid-like properties of the protein, while lack thereof indicates a solid-like state of the protein a) eGFP-NT2RepCT - show liquid-like properties of the protein condensates at pH 8, solid-like at pH 6. b) KSI-eGFP – exhibit solid-like properties at both pH 8 and 6, c) eGFP – exhibit liquid-like properties at both pH 8 and 6.

On the other hand, the inclusion bodies formed by KSI-eGFP (Figure 3b), remained intact upon cell damage at both pH of 6.0 and 8.0, respectively. In contrast, a spreading of the fluorescence signal from damaged cells expressing soluble eGFP was observed at both pH values (Figure 3c). As such these proteins can serve as effective reference points for describing the intracellular behavior of the biomimetic silk protein since their viscoelastic properties are not affected by the pH.

To further investigate the effect of the pH on the material state of the intracellular protein structures we carried out complete cell lysis in a buffer containing lysozyme and a detergent that fully disrupted cell structure (Figure 4). This treatment also allowed us to study the material state of the protein in the absence of the intracellular components known to have a crowding effect promoting protein LLPS [42].

**Figure 4.**
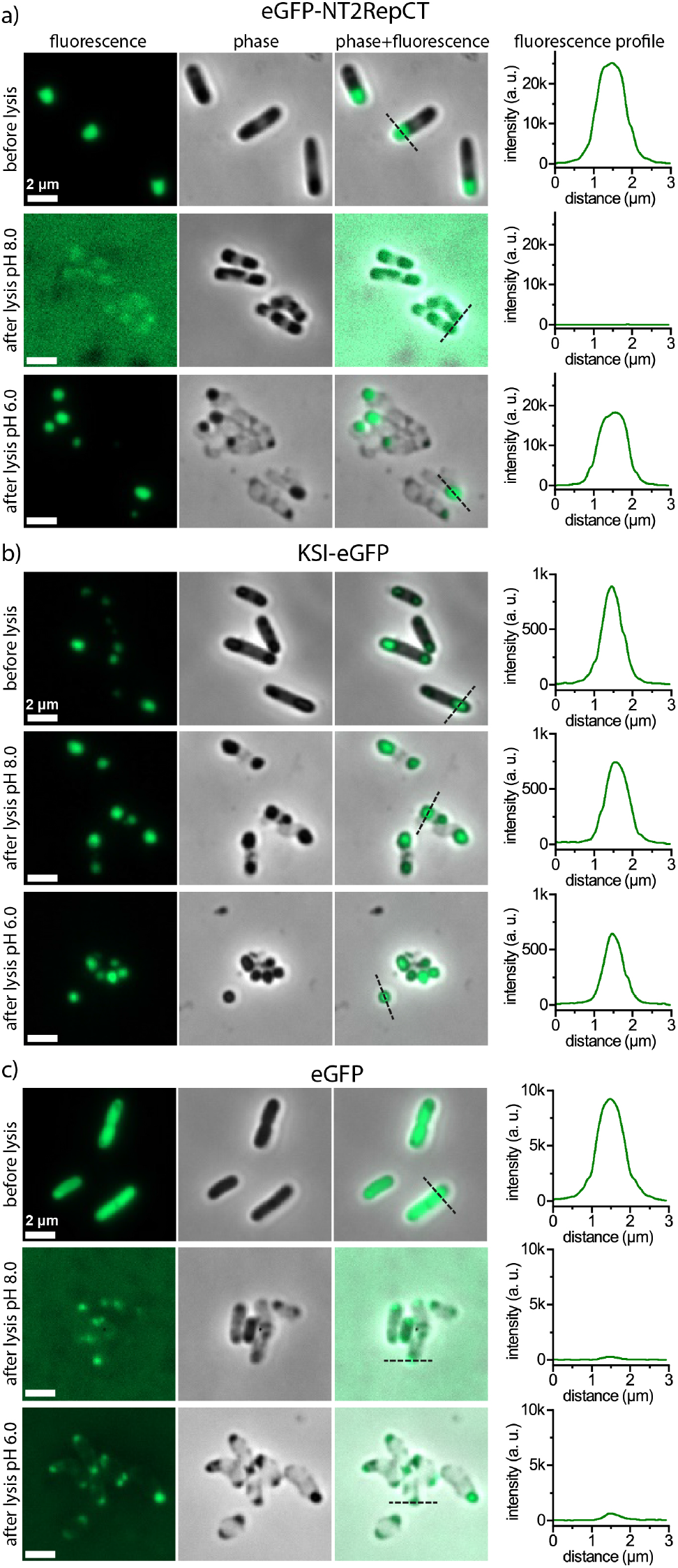
Comparison of fluorescence microscopy images of *E. coli* cells after recombinant expression of proteins under study before and after chemical lysis at pH 6.0 and pH 8.0. Graphs show fluorescence profiles - the intensity of the fluorescence signal as a function of the position on the dotted line displayed in the panel with the overlaid fluorescence and phase contrast channels. a) eGFP-NT2RepCT –show liquid-like properties of the protein condensates at pH 8 and solid-like at pH 6, b) KSI-eGFP – exhibit solid-like properties at pH 8 and 6, c) eGFP – exhibit liquid-like properties at pH 8 and 6.

The cells were imaged before and after lysis and the fluorescence signals and morphologies compared. In line with the previous experiment, we observed that the condensates formed by eGFP-NT2RepCT fully spread into the surrounding buffer at pH 8.0 but remained intact inside the cells at pH 6.0 (Figure 4a). An SDS-PAGE gel of the soluble and insoluble fractions of the crude lysed cells was performed further confirming that at pH 8.0 the protein remained soluble while at pH 6.0 the protein was mostly insoluble (Figure S1). In contrast, the intracellular structures formed by KSI-eGFP remained unaffected by the cell lysis and no spreading of the fluorescent protein was observed at either pH values(Figure 4b). Conversely, the fluorescence from the lysed cells expressing eGFP spread regardless of the pH indicating that the protein was soluble (Figure 4c). These results were also in line with the analysis of the cell lysates by SDS-PAGE showing that after cell lysis eGFP was present mainly in the soluble fraction at pH 6 and 8, while the KSI-eGFP was present only in the insoluble fraction at both pH values (Figure S1).

In addition, comparison of the fluorescence intensity acquired from the cross-sections of the cells before and after lysis provided quantification of the spreading of eGFP-labelled proteins. The comparison of these fluorescence profiles further confirmed the observations about the material state of the proteins. In the case of eGFP (the reference for soluble protein), the signal decreased by over 90 % to reach near baseline levels at both pH 6.0 and pH 8.0 (Figure 4c) while the signal for eGFP-KSI (the reference for insoluble protein), only decreased by approximately 20 % at both pH (Figure 4b). In contrast, the fluorescence profile of the cells expressing the biomimetic silk eGFP-NT2RepCT was pH dependent (Figure 4a). At pH 8.0, the intensity profile had decreased to near-baseline levels as observed in the cells expressing eGFP. At pH 6.0, however, the intensity profile decreased only by approximately 20 % in a similar way as what was observed in the case of eGFP-KSI.

### 3.3. Molecular crowding facilitates LLPS of NT2RepCT but does not affect its liquid to solid transition

Next, we investigated the LLPS properties of NT2RepCT (without eGFP fusion) *in vitro* - after purification, to serve as a comparison to its behavior observed *in vivo* (Figure 1-4). However, this comparison is not a straightforward one since the bacterial cytosol is composed of densely packed macromolecules placed in a confined space of the cell. In this highly crowded microenvironment biochemical processes including protein LLPS and aggregation are strongly affected by volume exclusion. Thus, to study the effect of molecular crowding *in vitro* a solution of a polymeric crowding agent (10 % dextran) was used to mimic the intracellular conditions that are present inside *E. coli* [42, 43].

The phase behavior of purified NT2RepCT was studied by phase contrast microscopy at different pH values. We observed the protein undergo LLPS forming condensates exhibiting liquid-like viscoelastic properties when it was mixed with buffers containing the crowding agent at a pH between 6.5 and 7.5 (Figure 5a, supplementary movie 1). Under these conditions, phase separation occurred at the lowest protein concentration of 1 mg/mL (30 *μ*M) (Figure S2). Next, we mixed the protein with a buffer at pH 6.0 and observed that it formed insoluble aggregates (Figure 5a – panel pH 6.0). Addition of the protein to more acidic buffers led to the clustering of protein aggregates (Figure 5a – panel pH 5.5) and formation of fibrillar aggregates (Figure 5a – panel pH 5.0).

**Figure 5.**
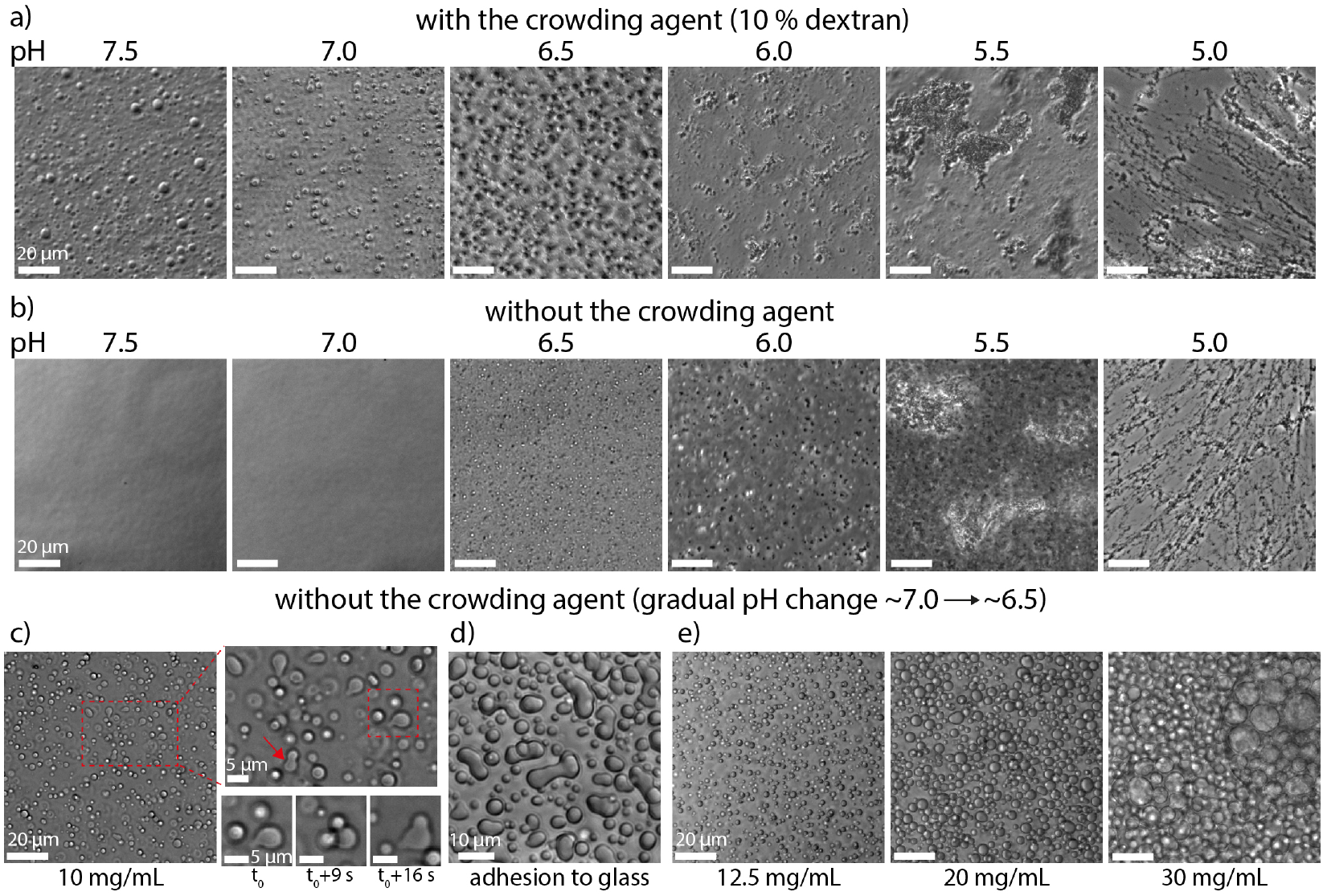
Phase transition of NT2RepCT *in vitro* studied by phase contrast microscopy at different pH values. In the presence of a molecular crowding agent (10 % dextran) and b) without the crowding agent (protein concentration 5 mg/mL) c) Images obtained during slow pH change from 7.0 to 6.5 (without dextran) showing liquid-like viscoelastic properties of protein microdroplets. High magnification and time-lapse images show fusion of the microdroplets. d) Adhesion and spreading of the microdroplets on a glass surface indicate their low interfacial tension. e) Increase of the protein concentration led to formation of larger microdroplets that merged faster.

Subsequently, the phase separating behavior of the purified NT2RepCT protein was investigated in the same pH range but without the addition of dextran as a crowding agent. The study was conducted to better understand the properties of the protein under conditions relevant to the previously established biomimetic silk fiber spinning process [24]. We observed that the protein did not undergo LLPS at pH 7.5 and 7 (Figure 5b) at a protein concentration of 5 mg/mL. LLPS occurred at pH 6.5 (Figure 5b) but the size of the condensates was smaller compared to the ones formed the presence of dextran, although the same protein concentration was used (Figure 5b vs 5a, panels at pH 6.5). At more acidic conditions the same behavior was observed as in the presence of the crowding agent. Regardless of the presence of crowing agent, the protein formed solid irregular aggregates at pH 6.0 or 5.5 (Figure 5b vs 5a – panels pH 6.0 and 5.5) and fibrillar-like aggregates at pH 5.0 (Figure 5b vs 5a – panel pH 5.0).

In order to verify the liquid-like properties of condensates at pH 6.5 without the crowding agent, we repeated the experiment using a higher protein concentration (10 mg/mL) (Figure 5c). Under these conditions, we observed merging of the droplets (Figure 5c – high magnification and time-lapse images) and adhesion and spreading of the droplets on a glass surface (Figure 5d). Furthermore, as protein concentration was further increased, larger droplets formed, and their fusion was readily observed (Figure 5e). This behavior is typical for liquid-like protein droplets that fuse into bigger spheres to minimize their surface tension [37].

Finally, the *in vitro* studies with and without molecular crowding agent were repeated with the purified eGFP-NT2RepCT to test a possible influence of the eGFP fluorescence tag on phase behavior of the protein. The results showed that the properties of the eGFP-NT2RepCT were the same as those observed for the protein variant without the fluorescent fusion partner (Figure S3 and Supplementary movie 2).

Overall, the *in vitro* studies (Figure 5) showed that in the presence of the crowding agent the purified NT2RepCT was able to undergo LLPS in a broader pH range (6.5 – 7.5) compared to the conditions without the crowing agent (in which LLPS was observed only at pH 6.5). This finding is in line with our *in vivo* studies showing that intracellular condensates of eGFP-NT2RepCT had liquid-like viscoelastic properties (Figure 2-3). Indeed, the pH of the cytoplasm of *E. coli* is in the range of 7.2 – 7.8 [44, 45]. Overall, the presence of the crowding agent did not alter the protein’s ability to undergo a liquid to solid transition that occurred in the same way as in the absence of the crowing agent (Figure 5a,b). These observations suggest that *in vitro* studies of the protein in the presence of a crowding agent can be linked with the assessment of the material state of the protein condensates *in vivo*.

### 3.4. NT2RepCT undergoes LLPS forming condensates that assemble into a continuous fiber in the presence of shear force applied during extrusion

Finally, we investigated the properties of the protein during a biomimetic extrusion process allowing the production of silk fibers. The protein solution was prepared in 20 mM Tris-HCl buffer, pH 8.0, as in the previous experiment (Figure 5,) and was extruded under constant pressure through a thin capillary tube (with an outlet diameter of approximately 20 *μ*m) which was immersed in a pH 5 coagulation buffer (500 mM sodium acetate buffer containing 200 mM NaCl). A similar use of pH-induced coagulation has previously been utilized for biomimetic spider silk spinning [24]. However, in our setup the outlet of the capillary tube was placed under an optical microscope equipped with a camera that enabled observation of the behavior of the protein at increasing concentration during extrusion into the coagulation buffer (Figure 6a).

**Figure 6.**
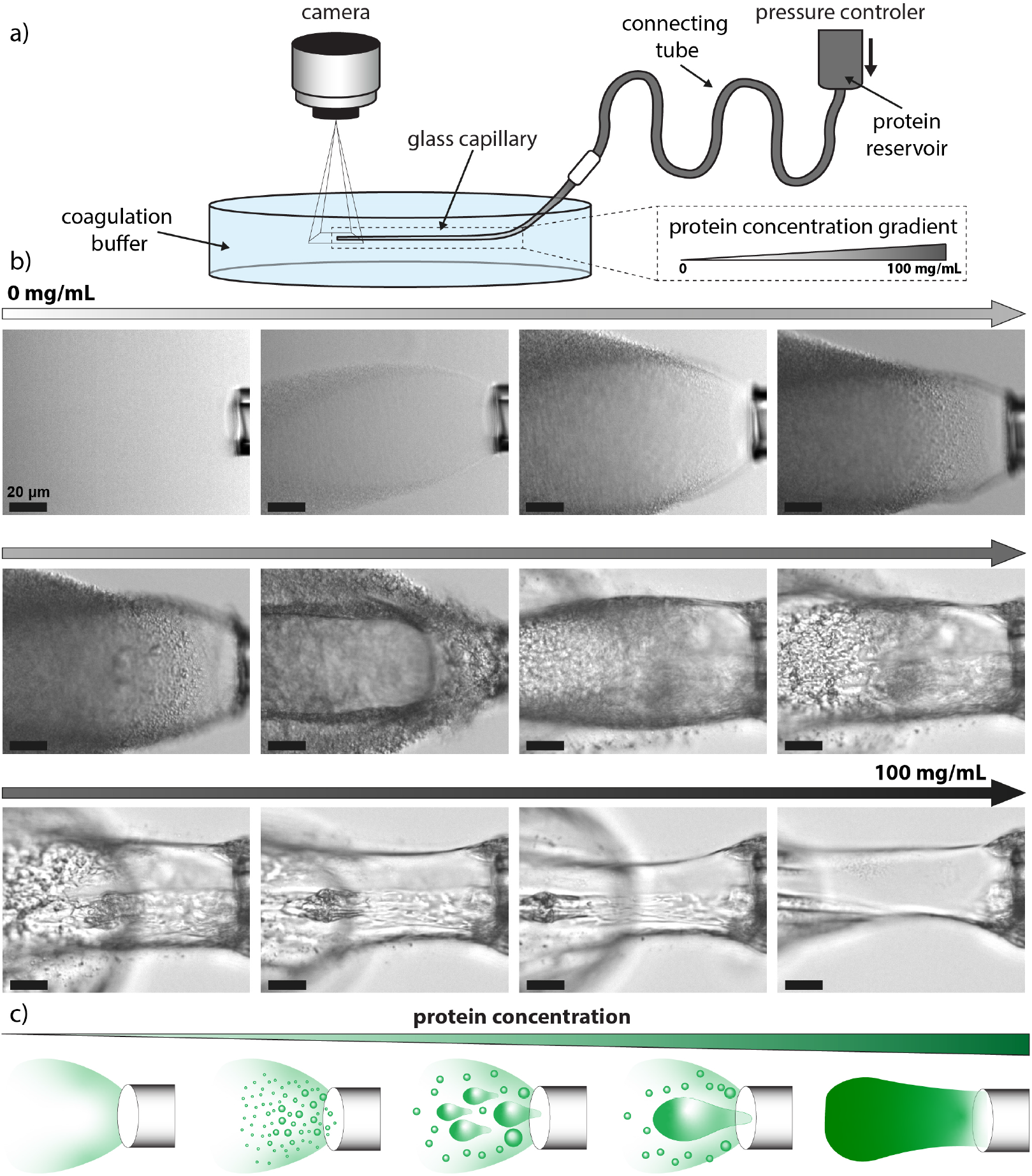
Investigation of the phase behavior of the NT2RepCT during extrusion through a capillary tube mimicking the process of biomimetic silk fiber spinning. a) schematic representation of the extrusion setup, b) time-lapse images of the extrusion of increasing protein gradient (0 - 100 mg/mL), full process presented in the Supplementary movie 3 and 4, c) schematic showing phase behavior of the NT2RepCT during the extrusion – at the lowest protein concentration LLPS is not observed, but an increase in the protein concentration leads to visible LLPS and formation of small protein droplets. With a further increase in protein concentration droplets grow in size, start to merge, and form a cohesive structure. At the highest protein concentration, no individual droplets are seen but a continuous fiber is formed.

To test whether LLPS of the protein occurs also during extrusion, we formed a gradient of protein concentration in the capillary tube (from 0 to 100 mg/mL) and carried out the extrusion gradually increasing protein concentration. At the same time, we monitored the events at the interface between the protein solution and the low-pH buffer (Figure 6b). We observed that, initially at low concentration, the protein mixed with the low-pH buffer, no LLPS was detected, and no fiber was formed. When the protein concentration increased, LLPS occurred and small protein droplets that dispersed in different directions could be observed. Further increase in the protein concentration led to the formation of bigger droplets. These, however, did not yet form a cohesive structure. Only subsequently, at the higher protein concentration, the droplets started to merge into a cohesive assembly. As the protein concentration increased further, a continuous fiber core started to form near the center of the capillary outlet. However, dispersed droplets could still be seen around the fibers’ core. Eventually when the protein reached a sufficiently high concentration (100 mg/mL) it immediately formed a continuous fiber that appeared as one single solid-like protein phase entering the buffer. (Figure 6b and Supplementary Movie 3 and 4).

Together these observations showed that LLPS of NT2RepCT occurred during the transition of the homogenous soluble protein into the solid-like fiber in a biomimetic extrusion process. We interpret this observation as suggesting that acidification triggers protein LLPS that leads to the formation of liquid-like condensates that rapidly shift to a solid-like state. At a high enough concentration, the shear force enables the formation of a continuous silk fiber. The intermediate stages in which only droplets were seen forming were only observed for protein concentrations lower than 100 mg/mL – well below the concentration required to form a continuous fiber (Figure 6c) [24]. Overall, our findings are in agreement with previously published studies showing LLPS of purified recombinant spider silk-like proteins and suggesting that liquid-like condensates are an intermediate protein state during fiber formation [11, 12].

## 4. Conclusions

Protein-based materials in Nature are formed through controlled protein condensation initiated by LLPS. Although many sequences of native proteins that assemble to build biological materials have been identified and their LLPS processes reproduced and studied *in vitro* using recombinantly produced proteins, the biotechnological strategies for the fabrication of bioinspired protein-based materials are still very limited. Currently, the major challenges are related to the identification of protein variants with an ability to undergo LLPS while at the same time exhibiting a high recombinant production yield without aggregating prematurely in their recombinant host and after purification. Our study shows that these criteria can potentially be screened for by utilizing recombinant expression in *E. coli* not only as a means of producing the proteins, but also to assess whether an overexpressed protein has the ability to undergo LLPS and form condensates with properties amenable for the engineering of protein-based materials.

Using the NT2RepCT biomimetic silk-like protein as an example, we demonstrated that intracellular LLPS of an overexpressed protein (a building block of a biological material) leads to its accumulation in a condensed liquid-like structure within the bacterial cell. In the case of NT2RepCT, its intracellular condensation is directly related to its function as a building block of silk fibers. The properties of the intracellular condensates the protein forms are similar to those of the spinning dope of native silk which enables storage of the highly concentrated protein in a liquid-like state in a pH range between 7.2 and 7.8 [44, 45, 46]. Once exposed to an acidic environment the material state of the protein condensates shifts from liquid-like to solid-like, a feature also found in native silk proteins [26, 41].

Intriguingly, KSI-eGFP (the reference protein known to accumulate in highly insoluble inclusion bodies), formed small, condensed structures early during overexpression that ultimately accumulated into solid-like assemblies. The superficial similarity of the initial part of the process as the protein localizes within the cell, may suggest that KSI-eGFP also initially undergoes LLPS. However, the protein would further aggregate into a more solid form in the molecularly crowded environment of the bacterial cytoplasm.

On the other hand, in the case of NT2RepCT, intracellular molecular crowding facilitated its LLPS and did not influence its ability to undergo the liquid to solid transition triggered by a decrease in pH. These properties could be reproduced using purified protein with the presence of a molecular crowding agent, showing that behavior of the protein *in vitro* can be related with its behavior *in vivo*. Moreover, LLPS of NT2RepCT occurred also during the process of its liquid to solid transition during biomimetic extrusion. However, liquid-like protein droplets could only be visualized when the protein concentration was too low to form a continuous fiber. This gradual transition from droplets to fiber suggests that at higher concentrations liquid-like condensates are still present, but they immediately fuse into a single-phase maturing into a continuous solid fiber. This therefore suggests that LLPS may play an important role influencing the final properties of the silk fibers, and thus, provides a link between the investigation of the intracellular phase behavior of a protein *in vivo* with its ability to form functional materials *in vitro*.

It is interesting that LLPS dominates the behavior of NT2RepCT so strongly that this behavior is also clearly seen already in the cell during protein synthesis. The question that follows is whether this connection is more generally applicable. If so, the protein’s phase behavior *in vivo* could be used as a strategy to help design proteins for protein-based materials. This could create new possibilities for bioinspired material engineering, for example enabling the use of *E. coli* as a “micro-test tube” in which characterization experiments could be done when screening for new material-forming proteins or when screening for variants with altered properties. The use of *in vivo* characterization would eliminate the need for purifying and characterizing each variant separately, thus speeding up the screening process. The tendency for LLPS or an overly high tendency for aggregation or solidification could be easily detected by such a screen. Even more refined characterizations can be conducted as our understanding of LLPS in material formation and *in vivo* processes develops.

## 5. Supporting Information

Figures S1-S3: SDS-PAGE of cell lysates after end point of recombinant expression, additional phase contrast microscopy images. Supplementary movies 1-4: LLPS of NT2RepCT and eGFP-NT2RepCT in a presence of the crowding agent, extrusion of NT2RepCT.

## 6. Acknowledgements

This work was carried out in the CatBat project funded by Novo Nordisk Foundation (#0061306) and under the Academy of Finland Center of Excellence Program (2022-2029) in Life-Inspired Hybrid Materials (LIBER), project numbers: 346105, 346112, 315140. In addition, we acknowledge Tiina Pessa-Morikawa form the HiLife Flow Cytometry Unit at Helsinki University for her help in conducting the flow cytometry analysis, and Ekaterina Osmekhina from Aalto University for help with designing the schematics used in Figure 6.

