## Supporting Information for "Recombinant protein condensation inside E. coli enables the development of building blocks for bioinspired materials engineering – biomimetic spider silk protein as a case study"

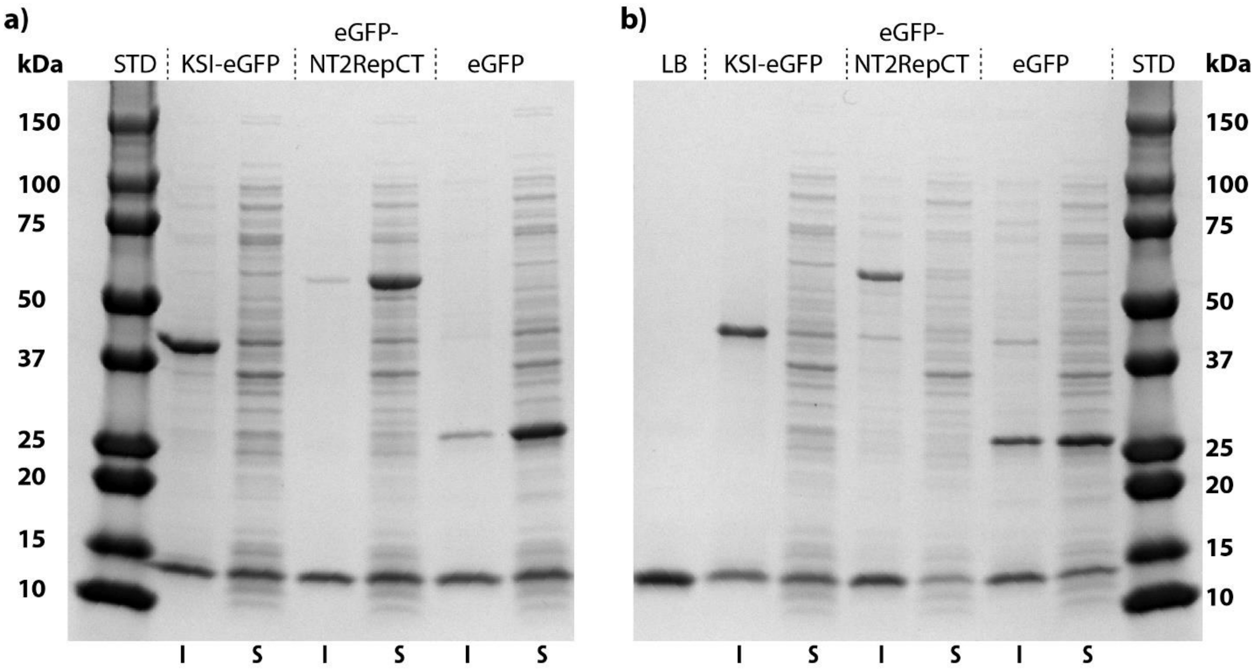


**Figure S1**. SDS-PAGE of cell lysates after end point of recombinant expression. **a)** at pH 8.0 and **b)** at pH 6.0. STD – molecular weight standard, I – insoluble fraction, S – soluble fraction, LB – lysis buffer. Predicted molecular weights: KSI-eGFP = 41.805 kDa, eGFP-NT2RepCT = 60.503 kDa, eGFP = 27.996 kDa, lysozyme (Sigma-Aldrich L7651) = 14.3 kDa (present in the lysis buffer).

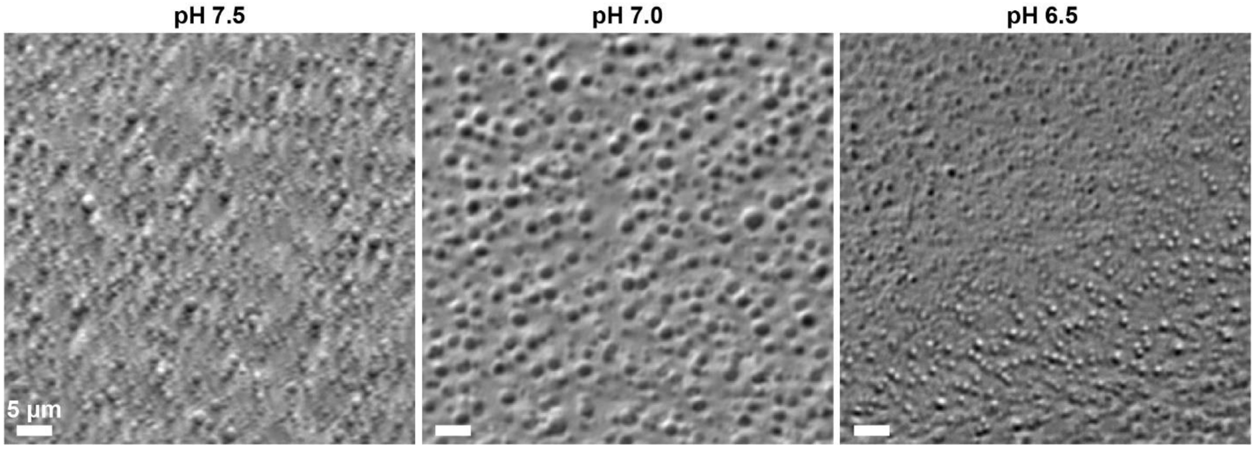


**Figure S2**. LLPS of NT2RepCT at different pH values in the presence of crowding agent (10 % dextran) at minimal protein concentration required for LLPS to occur (1 mg/mL).

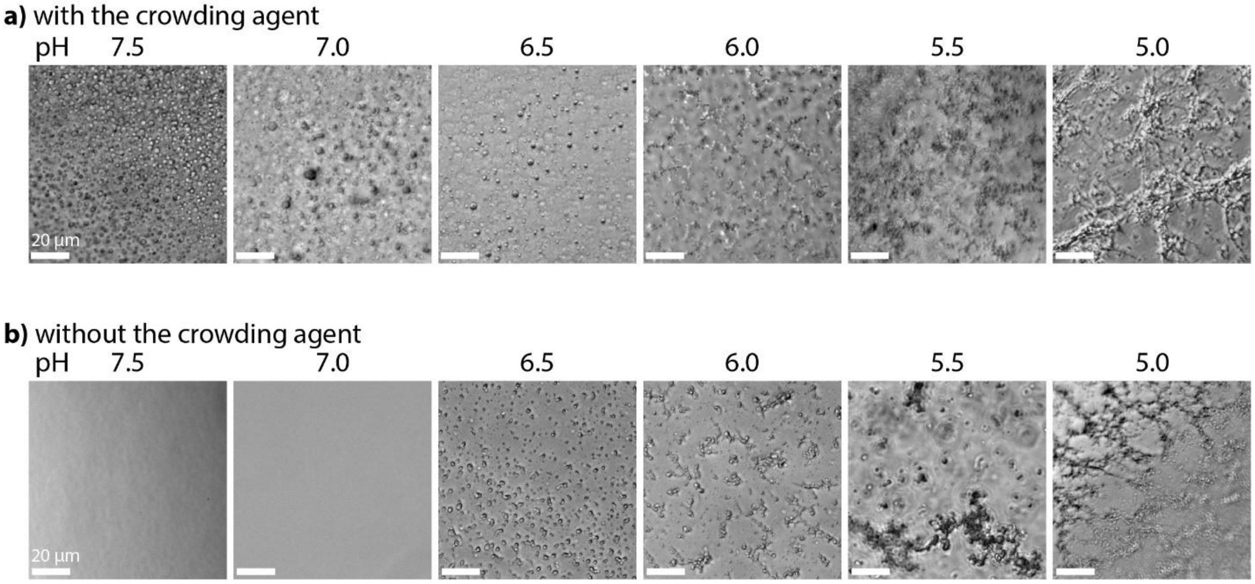


**Figure S3**. Phase transition of eGFP-NT2RepCT in vitro studied by phase contrast microscopy at different pH values. **a)** In the presence of a molecular crowding agent (10 % dextran) and **b)** without the crowding agent. Overall, these results show that LLPS properties of eGFP-NT2RepCT are consistent with the properties observed for the protein variant without the fluorescent fusion partner (Figure 5).

**Supplementary movies**:

**Movie 1** – LLPS of NT2RepCT in a presence of the crowding agent (10 % dextran) at pH 7.0, protein concentration 10 mg/mL. Fusion of protein droplets indicate their liquid-like viscoelastic properties.

**Movie 2** – LLPS of eGFP-NT2RepCT in a presence of the crowding agent (10 % dextran) at pH 6.5, protein concentration 10 mg/mL. Fusion of protein droplets indicate their liquid-like viscoelastic properties.

**Movie 3** – extrusion of increasing concentration gradient (0 to 100 mg/mL) of NT2RepCT through a capillary tube into low pH coagulation buffer.

**Movie 4** – extrusion NT2RepCT (100 mg/mL) through a capillary tube into low pH coagulation buffer.
